# The genetic basis for adaptation in giant sea anemones to their symbiosis with anemonefish and *Symbiodiniaceae*

**DOI:** 10.1101/2022.09.25.509434

**Authors:** Agneesh Barua, Rio Kashimoto, Konstantin Khalturin, Noriyuki Satoh, Vincent Laudet

## Abstract

Sea anemones in the order Anthozoa play an integral part in marine ecosystems by providing refuge and habitat for various organisms. Despite this, much of their molecular ecology remains elusive. Sea anemones can nurture numerous symbiotic relationships; the most iconic being the one between giant sea anemones and anemonefish. However, the genes and biological processes associated with this symbiosis in the sea anemones in unknown. Additionally, it is unclear how genes can mediate interactions between sea anemones, anemonefish, and symbionts from the algal family *Symbiodiniaceae*. Here we compared the gene expression profiles of tentacles from several cnidarians to uncover the genetic basis for adaptations in giant sea anemones to their symbiosis with anemonefish and *Symbiodiniaceae*. We found that tentacle transcriptomes of cnidarians are highly diverse, with closely related species having more similar expression patterns. However, despite an overall high correlation between gene expression and phylogeny, the giant sea anemones showed distinct expression patterns. The giant sea anemones had gene co-expression clusters enriched for processes involved in nutrient exchange and metabolism. These genes were not only differentially expressed, but also experienced evolutionary shifts in expression in giant sea anemones. Using a phylogenetic multilevel model, we found that *Symbiodiniaceae* and anemonefish significantly affect gene expression in giant sea anemone tentacles. By characterizing gene expression patterns, we identify genes and biological processes that provide evidence for the cross-talk between *Symbiodiniaceae*, anemonefish, and giant sea anemones. Our study demonstrates how integrated biological processes can lead to the evolution of a successful multi-organism interaction.

## Introduction

Anthozoans are a class of cnidarians that are the cornerstone of diversity in shallow and deep aquatic ecosystems (1). Several key innovations like modular, colonial growth forms; hard calcareous skeleton; and symbiosis with photosynthetic dinoflagellates (*Symbiodiniaceae*) have led to their ecological success. Due to their position as biodiversity hotspots and endangered status, coral reefs and their coral species are extensively studied (2, 3). However, several ecologically important anthozoans do not form reefs or have a hard calcareous skeleton, as exemplified by sea anemones (4). Sea anemones are soft-bodied, primarily solitary, and exclusively marine. They have evolved several unique adaptations, such as secretions that allow them to attach firmly to surfaces and venom systems that aid in prey capture and protection (5, 6). The starlet sea anemone *Nematostella vectensis* is now an established model system in Evo/Devo and has provided a treasure trove of knowledge into axial patterning mechanisms, nervous system development, and the evolution of fundamental bilaterian features (7). Although studies have provided deep insight into the above aspects of sea anemones’ biology, much of their molecular ecology remains elusive.

Tentacles are the primary tissues that sea anemones use to interact with their environment, with several species showing specialized tentacle morphologies (8, 9).

Studies of sea anemone tentacles have shed light on crucial evolutionary processes such as the origins of tissue specificity and cross-tissue recruitment of gene families (10, 11). Some sea anemone tentacles also house symbionts from the algal family *Symbiodiniaceae* (12, 13). The importance of symbiosis with *Symbiodiniaceae* has been studied extensively in corals, where photosynthetic products supplied by the symbiont assist the coral host in growth, metabolisms, reproduction, and survival (14, 15). Despite the importance of *Symbiodiniaceae* in corals, not all sea anemones form a symbiotic relationship with dinoflagellates (e.g. *Nematostella vectensis* and *Actinia tenebrosa*). In addition to *Symbiodiniaceae*, sea anemones form symbiotic relationships with other larger organisms such as fish or crustaceans (16–18). The most iconic of these relationships is the mutualistic symbiosis between anemonefish (*Amphiprion*; Pomacentridae) and giant sea anemone (19).

Ten species of giant sea anemones belonging to three different clades form a close mutualistic relationship with anemonefish (20–22). In this mutualistic interaction, the fish are sheltered by the venomous tentacles of the host sea anemone, and the fish provide the sea anemone with a source of nutrition (23, 24). Anemonefish are highly territorial and protect their host anemone from predators such as butterflyfish (*Chaetodon fasciatus*), thereby enhancing their survival (24, 25). Although there is a tight relationship between the fish and their host anemones, much of our information about this symbiosis comes from the perspective of the anemonefish, with little understanding of the molecular and evolutionary adaptations in the host anemones (20, 21).

In this study, we compare transcriptomes of tentacles from giant sea anemones with different species of cnidarians to uncover the genetic basis behind the unique ecology of anemonefish hosting giant sea anemones. The giant sea anemone showed gene expression patterns distinct from other cnidarians. Using a co-expression analysis, we identified groups of upregulated genes and processes related to symbiosis with anemonefish and *Symbiodiniaceae*. We identified genes that have experienced evolutionary shifts in gene expression in anemonefish hosting anemone using phylogenetic comparative methods.

Lastly, we tested for the effect of the relationship with anemonefish and *Symbiodiniaceae* on tentacle gene expression and found evidence of a potential adaptive effect. Our study identified important genes and biological processes involved in the multi-organism interactions between anemonefish, *Symbiodiniaceae*, and sea anemones and describes how the evolution of these genes and processes contributed to the formation of a successful multi-organism symbiotic relationship.

## Results

Anthozoans have similar tentacle gene expression patterns

To identify gene expression patterns of anthozoan tentacles, we analyzed publicly available RNA-seq data and generated data for this study (Table S1). Our sampled taxa comprised fourteen anthozoans, and as outgroup taxa, we included jellyfish (four scyphozoans, one cubozoa) and *Hydra* (hydrozoa) (Fig 1A). We included these taxa because they had publicly available data for tentacle gene expression. Along with gene expression data from tentacles, we also included data from tissues such as mesenteries, body column, and nematosomes. Most of our sampled taxa do not have a reference genome; therefore, we assembled *de novo* transcriptomes of all taxa for consistency. Mean BUSCO completeness was 82%, with jellyfish libraries having the lowest values (Table S2). The jellyfish libraries were of relatively poorer quality than the other taxa. Jellyfish tentacles are also highly heterogeneous in their gene expression, and the lower BUSCO values were not entirely unexpected (26). Our expression matrix comprised 1486 orthologs expressed across all tissues in all our sampled taxa.

**Figure 1.**
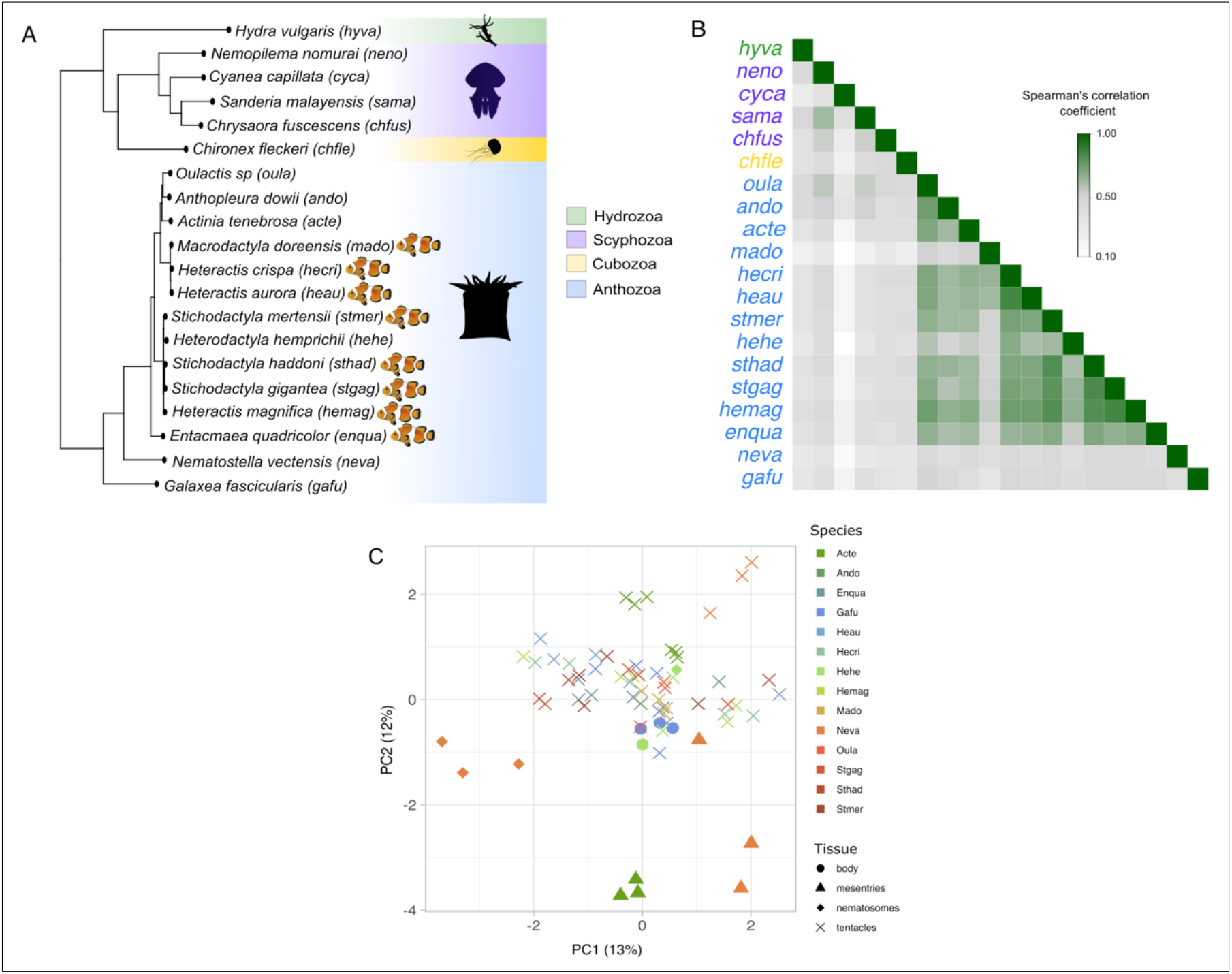
Tentacle gene expression is similar between closely related species. (A) Phylogeny of samples used in this study with species hosting anemonefish indicated. (B) Spearman correlations in tentacle gene expression between different cnidarians species. (C) Clustering of samples on the first two principal components showed a clear separation based on tissue. However, different tissues of *Heterodactyla* and *Galaxea* cluster with tentacles, suggesting that the gene expression in these tissues is similar to tentacles.

The variation in library quality could affect the ultimate gene count, which will affect the transcripts per million (TPM) quantification of gene expression. To make gene expression more comparable between incomplete references and organisms with varying gene counts, we used a modified metric called transcripts per million 10K (TPM10K) described in Munro *et al*. (27). TPM10K normalizes TPM to account for different sequencing depths and gene counts between species (see methods for details) (27).

We first checked for correlation (spearman rank correlation) between gene expression levels in our samples. Overall, gene expression within anthozoans was more similar than between anthozoans and non-anthozoans (Fig 1B, and Fig S1, S2, S3). However, there were some exceptions to this. For instance, *Galaxea fascicularis* (gafu) and *Nematostella vectensis* (neva) had expression patterns that were not similar to other anthozoans (Fig 1B). The highest correlations were between the giant sea anemones of the *Heteractis, Stichodactyla, Macrodactyla* and *Entacmaea* genera. Three interesting patterns emerged within these giant sea anemones. First, gene expression of *Macrodactyla doreensis* (mado) was more similar to expression patterns in *Heteractis* than in *Stichodactyla*. Second, *Heteractis magnifica* (hemag) had expression patterns more similar to *Stichodactyla* than other *Heteractis* species. Lastly, the giant sea anemone *Heterodactyla hemprichii* showed a comparatively lower correlation with the other giant sea anemone.

Next, we performed a principal component analysis (PCA) to explore gene expression patterns between different lineages and tissues (Fig 1C). Since anthozoans show expression patterns very different from non-anthozoans, we discuss the results of PCA using only anthozoans. The PCA with all cnidarian samples is included in the supplementary information (Fig S4). The first two components partition the data based on tissues. This suggested that generally, tentacles have an expression pattern distinct from other tissues in sea anemones. However, non-tentacle tissues of *Galaxea fascicularis* and *Heterodactyla hemprichii* cluster together with tentacles, suggesting that the expression of these tissues is similar to that of tentacles.

The comparative transcriptome analyses showed that anthozoan tentacles have similar gene expression patterns and are distinct from non-tentacle tissues. Additionally, tentacle gene expressions of giant sea anemones, especially those hosting anemone fish, were highly correlated. The similarity of closely related lineages suggests a strong phylogenetic component to tentacle gene expression.

### Giant sea anemone tentacles evolved convergent gene expression patterns

The high similarity in gene expression between closely related taxa suggests a strong phylogenetic component. To better understand the relationship between gene expression and evolutionary history, we compared sequence-based-phylogenetic distances with expression distances between all species pairs. To obtain the phylogenetic distances, we constructed a phylogenetic tree using one-to-one single-copy orthologs and estimated expression distances using 1-Spearman coefficients (pairwise). Mantel test between phylogenetic distance and expression distance matrices showed that they were significantly correlated (*R* = 0.651, p-value = 0.001) (Fig 2A). To get an idea of the magnitude of expression change with an increase in the phylogenetic distance, we fit a simple linear model to the distance data, which estimated a slope of 3.081 (adjusted *R2 =* 0.62, p-value < 0.0001) (Fig 2A). These results suggest that closely related taxa tend to have similar tentacle gene expression patterns, with expression variation increasing with increased phylogenetic distance.

**Figure 2.**
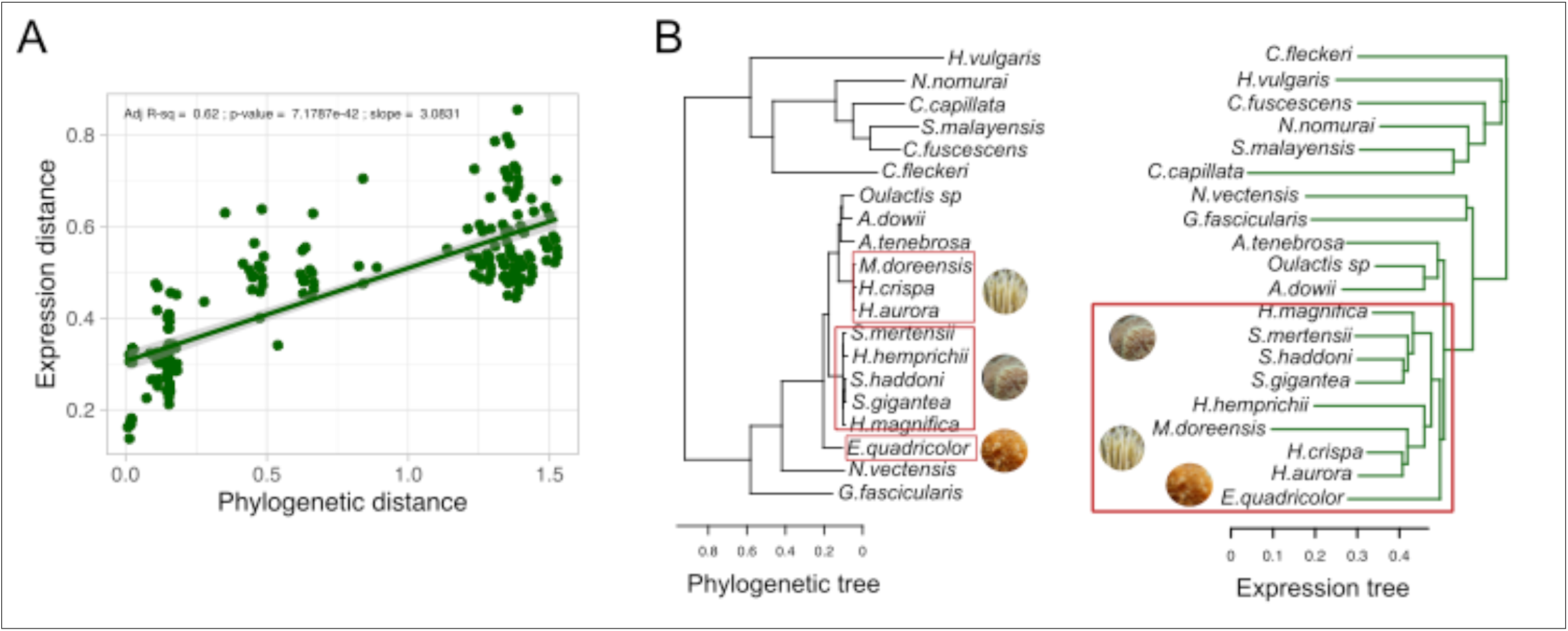
Although there is an overall consensus between expression and phylogeny, giant sea anemones have converged on a distinct tentacle gene expression pattern. (A) There is a significant positive relationship (R2 = 0.62) between phylogenetic distance and expression distance (1-Spearman coefficient) implying that expression divergence increases with phylogenetic distance. (B) There is an overall consensus between the phylogenetic tree and neighbor joining tree of expression distance. However, the giant sea anemones, which form phylogenetically separate clades, form a single monophyletic clade when looking at their expression (red boxes); suggesting a likely convergence of expression patterns. Images from top to bottom in phylogenetic tree: *H*.*crispa, S*.*gigantea, E*.*quadricolor*.

The high slope of the linear model (which can be interpreted as high evolutionary rates (28)) suggested that lineages may have evolved highly divergent tentacle gene expression patterns. We compared the species’ tree with the tentacle gene expression tree to determine whether gene expression patterns deviated from their expected evolutionary trajectory (Fig 2B). We constructed the expression tree using the 1-Spearman coefficient distances, while the species tree was constructed using a multisequence alignment of 1:1 single-copy orthologs using IQ-TREE 2 (29). The species tree had high bootstrap support and was consistent with previously estimated cnidarian phylogenies and an updated phylogeny of giant sea anemones (20–22, 30, 31). At a higher level, the species and expression trees were consistent; there was a clear separation of anthozoan and non-anthozoan lineages in both trees, *Nematostella vectensis* and *Galaxea fascicularis* formed divergent lineages (to giant and other sea anemones) in both the species and expression trees, and sea anemones *Actinia tenebrosa, Anthopleura dowi*i, and *Oulactis sp* were grouped with giant sea anemones. Despite the overall consistency, there were several key differences between the species and expression trees when looking at internal branches.

In the species tree, the different species of giant sea anemones formed separate lineages, consistent with previous results (20–22). However, instead of distinct lineages in the expression tree, they form a single large clade (Fig 2B). This suggests evolutionary forces have caused giant sea anemone tentacles to evolve convergent gene expression phenotypes.

### Genes upregulated in giant sea anemone tentacles are involved in the metabolism and biosynthesis of organic compounds

To get a sense of the biological underpinnings in giant sea anemone tentacles, we identified genes and processes that were differentially regulated. We performed differential gene expression (DGE) analysis using pairwise comparisons of *Heteractis, Stichodactyla, Macrodactyla*, and *Entacmaea* with *Actinia tenebrosa, Galaxea fascicularis, Nematostella vectensis*, and *Oulactis sp*. We identified several differentially expressed genes in the giant sea anemones. In total, 222 orthologous genes were upregulated in all giant sea anemones, while 240 orthologous genes were downregulated (Table S3 and Table S4). To determine the functional implications of these orthologs, we estimated co-expression clusters using the R package *coseq* (32). Among the upregulated genes, 222 were grouped into two clusters (cluster up-1 = 46 genes, cluster up-2 = 176 genes). 240 downregulated genes were also grouped into two clusters (cluster down-1 = 202 genes, cluster down-2 = 38). Genes were included in clusters based on *Maximum a Posteriori* scores, a Bayesian optimized parameter search approach for estimating joint probability distributions (33). The genes in cluster up-1 have expression patterns varying between samples (both within and between species). For example, genes in cluster up-1 tend to have an overall higher expression in *Entacmaea quadricolor*, while in *Heteractis magnifica* they tend to have a generally lower expression (boxplot in Fig 3A). In contrast, genes in cluster up-2 have a more uniform expression pattern between samples and species. The correlation plot for cluster up-2 shows patterns similar to the correlation plot in Fig 1; where closely related species have higher expression similarity, *Heteractis magnifica* has expression patterns similar to *Stichodactyla* than *Heteractis*, and *Macrodactyla doreensis* has expression patterns similar to *Heteractis* than *Stichodactyla*. Interestingly, it appears that *Macrodactyla doreensis* genes in cluster up-2 have an inverse relationship with *Stichodactyla* in cluster up-2 (grey tiles in correlation plot). This inverse relationship was also observed between *Heteractis magnifica* confirming that *Heteractis magnifica* has expression properties more similar to *Stichodactyla*. The variability in correlations in cluster up-1 could be due to noise generated by using a few gene samples. We estimated correlations using a random assortment of 47 genes to check if this was the case. A random set of 47 genes showed less noisy correlation patterns (Fig S5), suggesting that the 47 genes in cluster 1 are highly variable due to biological factors and not sampling bias.

**Figure 3.**
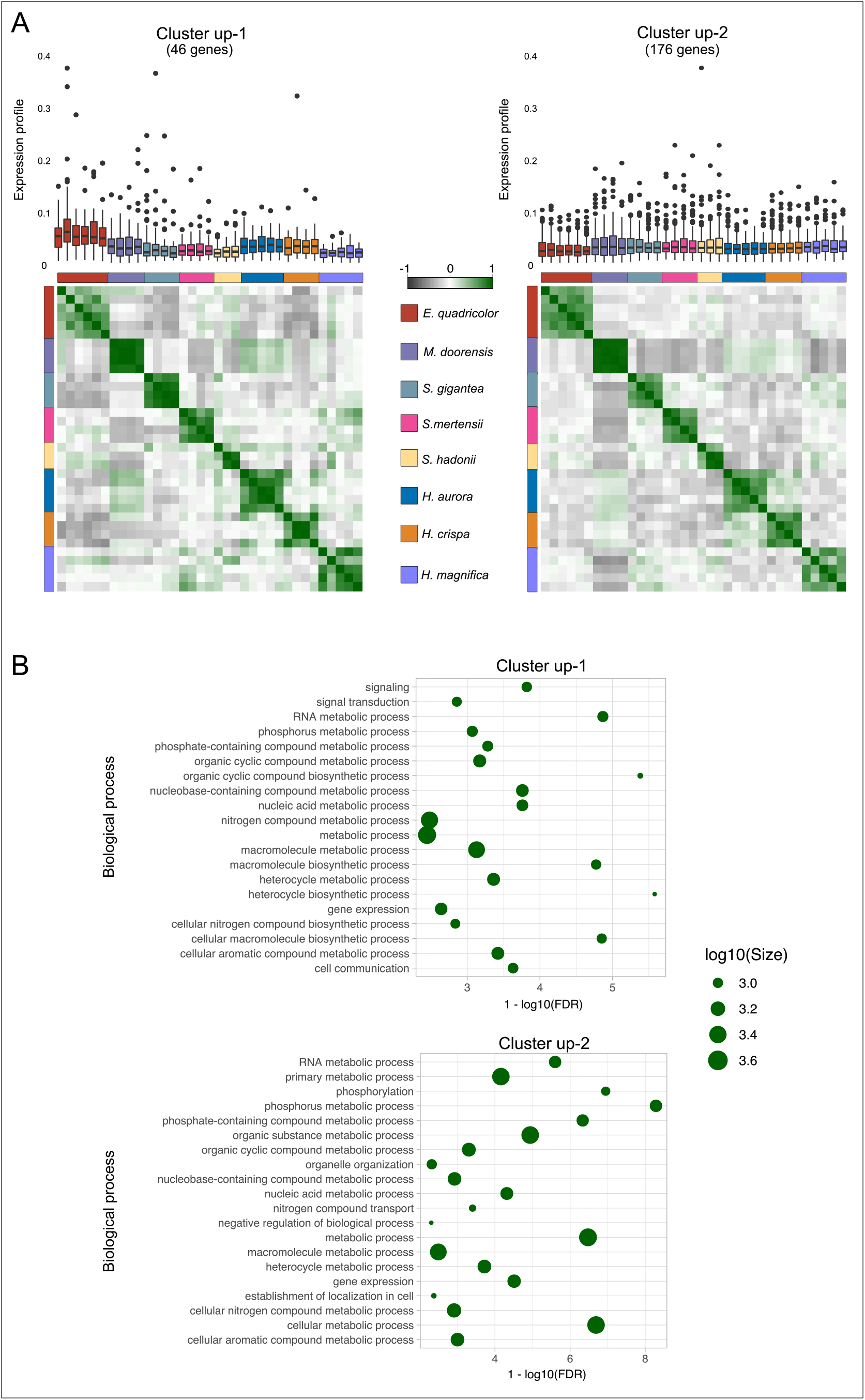
Groups of co-expressed genes that are upregulated in giant sea anemone are involved in processes relating to metabolism and biosynthesis of organic compounds, reflecting their functional relationship with *Symbiodiniaceae*. (A) Expression profiles show how expression of each co-expression cluster varies across different species of giant sea anemone. The boxplots represent mean expression profiles (with upper and lower quartiles) of cluster genes in each sample. The dots outside the bars indicate outliers. Despite many outliers, expression profiles are consistent across samples within species. Individuals of the same species have a high correlation in gene expression for cluster up- 2 (correlation plots). For cluster up-1, there is higher inter-species variation. For instance, different species of *Stichodactyla* and *Heteractis* have a relatively higher correlation for genes in cluster up-2 than in cluster up-1. Gene expression for cluster up-2 appears to be more lineage-specific, while gene expression for cluster up-1 is more varied. (B) Both clusters up-1 and up-2 are enriched for GO terms related to metabolisms and biosynthesis of organic compounds (only the top 20 are shown). Size of circles represents number of genes associated with the specific GO terms. Horizontal axis represents increasing significance of the association. *Symbiodiniaceae* are key in mediating the processes in both clusters and could be responsible for the species-specific expression pattern.

We annotated the genes from the upregulated clusters using Gene Ontology (GO) terms and performed an enrichment analysis to identify biological processes overrepresented in the two clusters. GO term enrichment showed that both clusters were overrepresented with terms related to biosynthesis and metabolic processes. Fig 3B shows the top 20 GO terms based on ontology size (for the complete list, see Table S5). In addition to the terms in Fig 3B, which represent general biological processes, several GO terms provided insight into more specific roles played by genes from these clusters. For example, cluster up-1 was enriched for the GO term GO:0044419: *biological process involved in interspecies interaction between organisms* (Table S5). The ontology neighborhood of this GO term was populated with terms that described symbiotic interactions and relationships with other organisms (34). Cluster up-2 was enriched for terms related to developmental processes such as GO:0007417- central nervous system development, GO:0060322- head development, and GO:0042461- photoreceptor development (Table S5). These fundamental biological processes might explain the more uniform gene expression patterns in cluster up-2 compared to cluster up-1.

The two clusters of downregulated genes showed similar overall trends to the upregulated clusters (Fig S6). Cluster down-1 replicated the lineage-specific expression pattern observed in cluster up-2 of the upregulated genes (Fig S6). In contrast, cluster down-2 of the downregulated genes was similar in expression variation to cluster up-1 of the upregulated genes. GO term enrichment of the downregulated clusters showed that they both comprised genes mainly involved in metabolic and catabolic processes (Fig S6 and Table S7, Table S8). Cluster down-1 also contained genes involved in fundamental processes like organelle organization and development, mirroring the expression pattern in cluster up-2 of the upregulated genes. The highly variable clusters of the upregulated and downregulated genes were enriched for different processes. The highly variable upregulated cluster (up-1) was enriched for processes related to biosynthesis. In contrast, the highly variable downregulated cluster (down-2) was enriched for catabolic processes. The differential gene expression and co-expression analyses provided evidence for crucial functional differences between giant sea anemones and other anthozoans. The giant sea anemones have higher expression of genes related to the metabolism and biosynthesis of nitrogenous compounds, phosphate-containing compounds, and other essential biomolecules (Fig 3B). These processes are highly relevant for symbiotic relationships between organisms where the exchange of nutrients occurs. The symbiotic relationship of giant sea anemones with anemonefish and *Symbiodiniaceae* could have influenced gene expression in the tentacles and contributed to the evolution of their unique gene expression profiles.

### Gene expression divergence and the relationship between anemonefish and *Symbiodiniaceae*

Giant sea anemones have a complex symbiotic relationship with *Symbiodiniaceae* and anemonefish. These symbiotic relationships could have influenced the evolution of gene expression patterns in giant sea anemones. We carried out a phylogenetic analysis of variance (phy-ANOVA) to determine whether genes in species hosting anemonefish have experienced evolutionary shifts in expression compared to non-hosting species. We also used a phylogenetic multilevel model (PGMM), to checked for the combined effect of hosting anemonefish and *Symbiodiniaceae* on gene expression.

We carried out the phy-ANOVA using the Expression Variance and Evolution (EVE) model developed by Rohlfs and Nielsen which aims to identify genes with high expression divergence as a result of adaptation in candidate species (35). We ran the EVE model using the clusters of upregulated and downregulated orthologs obtained from the co-expression analysis and branches leading to *Macrodactyla, Heteractis, Stichodactyla*, and *Entacmaea* as the test branches (Fig 4A). The EVE model found significant evidence (p- value < 0.05) of gene expression shifts in 99 upregulated genes and 9 downregulated genes (Table S5 and S6). To improve specificity, we focused on the ten genes with the highest magnitude of theta shift (labelled genes in Fig 4B top panel). Amongst the ten genes with the highest magnitude of theta shift, all of them were upregulated. These ten genes regulated biosynthetic and metabolic processes (Table S9).

**Figure 4.**
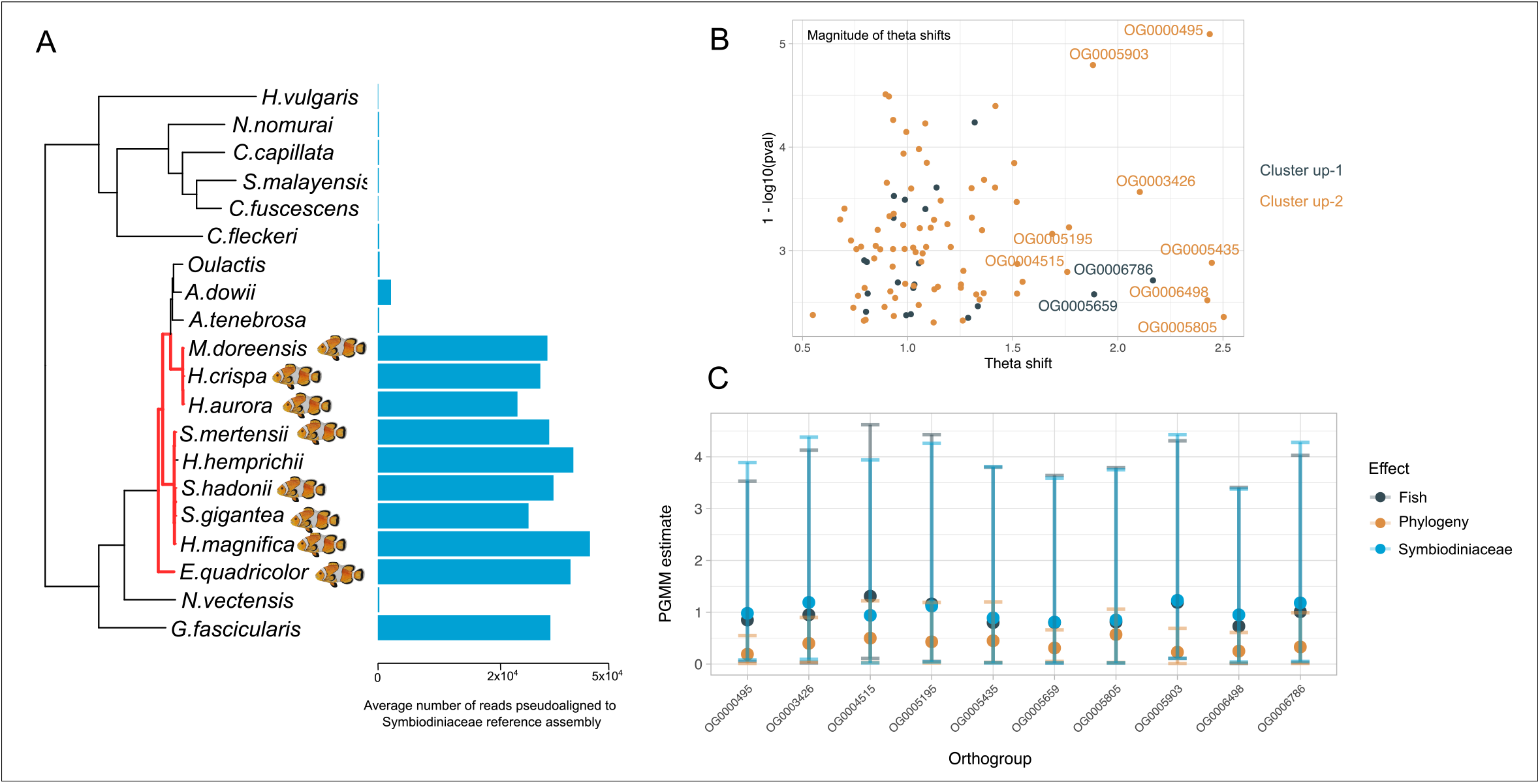
*Symbiodiniaceae* and anemone fish have influenced gene expression evolution in the tentacles of giant sea anemone. (A) Using the EVE model we tested whether lineages of giant sea anemone hosting anemone fish experienced significant shifts in gene expression evolution. Test branches are labelled red. Although *Heterodactyla hempirchii* is nested within the giant sea anemones, they do not host anemone fish, thus it was not included as test branch. The bar plots with the phylogeny show the average number of pseudoaligned reads to a composite *Symbiodiniaceae* reference transcriptome (see methods). These measurements were used to classify anemones as possessing or not possessing *Symbiodiniaceae*, which were used in the PGMM. (B) The panel shows genes that experienced significant shifts (EVE model theta shifts) in gene expression throughout evolution. The top 10 genes (labelled) were used to fit a PGMM to estimate the effect of phylogeny, *Symbiodiniaceae*, and anemone fish on tentacle gene expression. (C) The lower panel shows the estimates of the PGMM. Although the effect sizes are low and confidence intervals are wide, they do not overlap zero implying that the presence of anemone fish and *Symbiodiniaceae* have an effect on gene expression evolution in tentacles of the giant sea anemone.

We fit a PGMM using gene expression data from the ten genes with the highest gene expression shifts. The PGMM can estimate the effects of symbiosis with anemonefish and symbiosis with *Symbiodiniaceae* on gene expression evolution. To determine whether an anemone had symbiosis with *Symbiodiniaceae*, we checked for the abundance of *Symbiodiniaceae* reads in the assembled transcriptomes of our sampled cnidarians using a pseudoalignment approach (see methods). The bar plots adjacent to the phylogeny (Fig 4A) show the average number of reads that were pseudoaligned with *Symbiodiniaceae* reads. Our approach captured known trends in *Symbiodiniaceae* distribution, with the giant sea anemones, *Anthopluera*, and the coral *Galxea fascicularis* having the highest abundance of *Symbiodiniaceae* (36–38). Although a few reads mapped to sea anemones like *Actinia tenebrosa, Nematostella vectensis*, and jellyfishes, these were negligible and likely represented highly conserved transcripts. For our analysis, we classified organisms with these low abundant reads as not possessing *Symbiodiniaceae*. The modelling was done under a Bayesian framework using the *brms* R package (39). Mean expression levels of the ten genes were fit as multivariate response variables with the presence/absence of anemonefish and *Symbiodiniaceae* fit as the group-level effects (random effects in the terminology of mixed models). Phylogenetic relationships were modelled as a group-level effect to account for the non-independence between species. The parameter estimates for each effect (phylogeny, anemonefish, and *Symbiodiniaceae*) were low and with wide confidence intervals (95%-CI); however, the lower end of the CI did not overlap zero (Table S10), suggesting that all three parameters affected gene expression. The low effect sizes were likely due to low power as our sampling only included two lineages with *Symbiodiniaceae* and without anemonefish. Expanding the dataset to include more species of non-anemonefish hosting anemones possessing *Symbiodiniaceae* would provide better estimates and help tease out the relative effects of *Symbiodiniaceae* and anemonefish on gene expression variation. However, despite our limited dataset we still get evidence of a significant effect.

The EVE model and PGMM analysis showed that several genes in giant sea anemones had altered their expression throughout their evolutionary history, with symbiotic relationships with anemonefish and *Symbiodiniaceae* having a non-zero effect on gene expression evolution. The role of these genes can provide clues as to how specific biological processes can mediate interactions between organisms in a multi-organism symbiotic relationship.

## Discussion

We identified genes and biological processes relevant to multi-organism symbiosis and show how they experienced specific evolutionary trends in giant sea anemone lineages. In this section, we discuss how these genes and processes help mediate the symbiotic relationships between the anemones, anemonefish, and *Symbiodiniaceae*. We conclude by describing how the multi-organism interaction can promote the growth and ecological success of all the species involved.

### The role of gene co-expression clusters in multi-organism symbiosis and evolution of gene expression in giant sea anemone tentacles

Understanding the evolution of specific gene families and biological processes can provide insights into how anthozoans adapt to different ecological challenges (6, 40). We identified gene co-expression clusters in giant sea anemones primarily involved in the metabolism and biosynthesis of organic compounds. The heightened activity of these processes is the hallmark of the cnidarian-dinoflagellate symbiosis.

Studies using nanoscale secondary ion mass spectrometry and stable isotope labelling showed nutrient uptake and translocation at the organismal and cellular scales between host and symbiont (41). Using giant sea anemone (*H. crispa*) and mass spectroscopy, Verde *et al*. demonstrated a direct transfer of nitrogen and carbon-containing products from the host anemone to the endosymbiotic zooxanthellae and anemonefish (42). Nutrient exchange can also occur in the reverse direction between anemonefish to host anemone and endosymbiotic zooxanthellae (43). These studies reveal an active nutrient exchange between anemonefish, their host anemone, and *Symbiodiniaceae* (36). The gene and processes upregulated in the co-expression analysis likely mediate this nutrient exchange and help in the metabolism of biomolecules.

Genes exhibiting expression level variance between species can harbor adaptive variations that affect expression levels or could just be responding to environmental cues (35). We believe the gene expression variation observed in co-expression cluster up-1 is likely in response to genetic and environmental factors specific to each individual. In other words, the external environment can influence the expression of specific genes leading to differences in metabolic and biosynthetic processes in giant sea anemones tentacles. For instance, symbiont activity in *Anthopleura* is susceptible to external food availability, shifting the balance between heterotrophic and autotrophic lifestyles (44). Sea anemones absorb more ammonia during the daytime than at night, indicating that ammonia uptake is driven by the photosynthetic activity of *Symbiodiniaceae* (45). Furthermore, the presence of anemonefish also influences nitrogen uptake; anemonefish consume zooplankton during the day, following which they promptly excrete, providing the sea anemone and photosynthetically active *Symbiodiniaceae* with a rich source of nitrogen that they rapidly absorb (45). Therefore, differences in environmental conditions between species (and even individuals) of sea anemones will influence the activity of metabolic and biosynthetic processes in tentacles, leading to wide variation in gene expression.

From the EVE analysis, we identified several genes that show evidence for divergent gene expression in giant sea anemones. These genes were primarily involved in regulating metabolic and biosynthetic processes and developmental functions. While it is difficult to ascribe a specific function to a gene without molecular assays, we can still understand the role they potentially play by looking at how those genes work in other organisms. Here we discuss the roles of a few important ones. The list of genes and their annotations can be found in Table 1 and (Table S11).

**Table 1.**
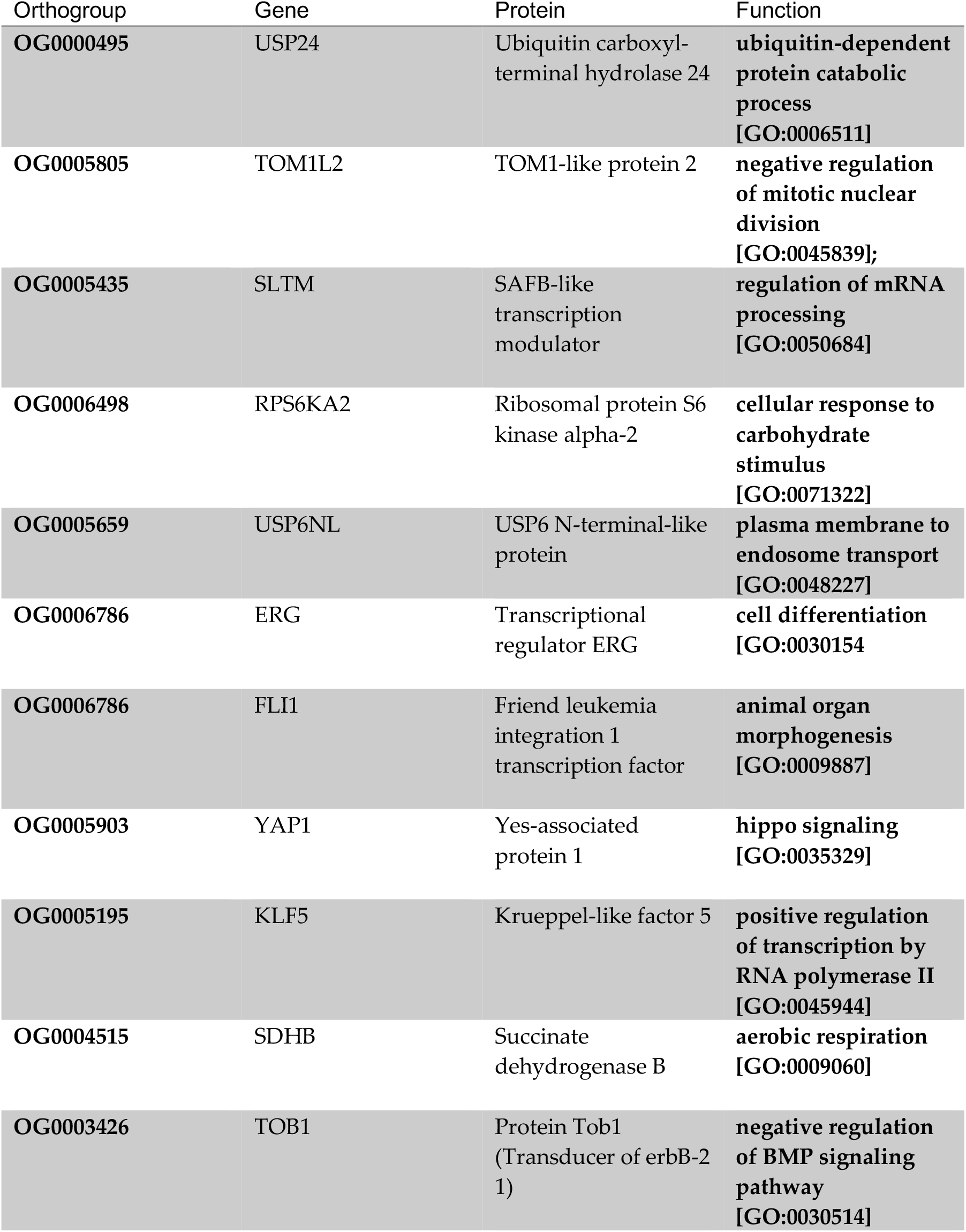
Top 10 genes with the highest rate shifts identified from the EVE analysis.

| Orthogroup | Gene | Protein | Function |
| --- | --- | --- | --- |
| OG0000495 | USP24 | Ubiquitin carboxyl-terminal hydrolase 24 | **ubiquitin-dependent protein catabolic process**<br>[GO:0006511] |
| OG0005805 | TOM1L2 | TOM1-like protein 2 | **negative regulation of mitotic nuclear division**<br>[GO:0045839]; |
| OG0005435 | SLTM | SAFB-like transcription modulator | **regulation of mRNA processing**<br>[GO:0050684] |
| OG0006498 | RPS6KA2 | Ribosomal protein S6 kinase alpha-2 | **cellular response to carbohydrate stimulus**<br>[GO:0071322] |
| OG0005659 | USP6NL | USP6 N-terminal-like protein | **plasma membrane to endosome transport**<br>[GO:0048227] |
| OG0006786 | ERG | Transcriptional regulator ERG | **cell differentiation**<br>[GO:0030154] |
| OG0006786 | FLI1 | Friend leukemia integration 1 transcription factor | **animal organ morphogenesis**<br>[GO:0009887] |
| OG0005903 | YAP1 | Yes-associated protein 1 | **hippo signaling**<br>[GO:0035329] |
| OG0005195 | KLF5 | Krueppel-like factor 5 | **positive regulation of transcription by RNA polymerase II**<br>[GO:0045944] |
| OG0004515 | SDHB | Succinate dehydrogenase B | **aerobic respiration**<br>[GO:0009060] |
| OG0003426 | TOB1 | Protein Tob1 (Transducer of erbB-2 1) | **negative regulation of BMP signaling pathway**<br>[GO:0030514] |

One of the genes with high expression divergence encodes Yes-associated protein (YAP), a transcriptional coactivator of the Hippo pathway involved in cell differentiation, stemness, and cell proliferation (46). The Hippo pathway is exceptionally conserved in metazoans and regulates cell growth and proliferation in a highly dynamic manner (46, 47). Several upstream signals like cell-cell contact, cell polarity, and soluble factors (hormones and growth factors) determine the activity of YAP and the Hippo pathway (46). Two basic modes of signaling are highly relevant in the context of giant sea anemones and their symbioses; the mechanical strain on cells and energy stress. Mechanical strain can cause stretching of cells which has been shown to increase the translocation of YAP into the nucleus resulting in increased cell proliferation (48). While in their host anemones, anemonefish routinely exhibit behaviors such as fanning and rapid fin strokes that cause turbulent water flow (49). The turbulence can lead to alternate periods of stretching and relief of mechanical strain in the cells of the giant sea anemone tentacles which could increase the activity of YAP, thereby promoting growth. Like other pathways involved in organ growth and tissue homeostasis, cellular stress can also perturb the Hippo pathway. There is a strong connection between glucose-mediated cellular metabolic status and the Hippo pathway, where glucose deprivation robustly inhibits YAP activity (46, 50). Glucose is a major metabolite transferred between *Symbiodiniaceae* and their cnidarian hosts, providing for much of the host’s total energy needs. (51, 52). Therefore, the symbiosis with *Symbiodiniaceae* ensures high glucose levels, which keeps YAP activity high, enhancing the growth potential of giant sea anemones.

The importance of YAP and the Hippo signaling pathway was well illustrated in a recent study in *Hydra (53)*. The researchers showed that the Hippo pathway regulates axis formation and morphogenesis in hydra, with YAP having a critical role in increasing cell proliferation, especially in the tentacles (53). The Hippo pathway and expression of YAP had important implications for the evolution and origin of axis formation in Metazoans (53). Similarly, the heightened activity of YAP and the Hippo pathway, along with the positive feedback effect of anemonefish and *Symbiodiniaceae*, might have enabled giant sea anemones to evolve their large size.

Another transcription modifier showing divergent gene expression in giant sea anemones was a member of the Kruppel-like factors (KLFs). KLFs are zinc finger proteins that can bind to regulatory elements on the DNA to either activate or repress transcription (54). In recent years, KLFs have emerged as a major metabolic regulator where they transcriptionally control critical processes in metabolism, nutrient uptake, and tissue utilization of macromolecules like carbohydrates, lipids, and amino acids (55). Another essential function of KLFs is their ability to respond to periods of nutrient stress and excess, thereby acting as a molecular switch, shifting metabolism between periods of nutrient storage and nutrient utilization (55). This strong association of KLFs with the cell’s metabolic state makes them ideal regulators for nutrient exchange between giant sea anemone tentacles and *Symbiodiniaceae*.

While KLF and YAP have potentially beneficial functions for multispecies interaction, several genes have functions suited to the general growth and development of the giant sea anemone. For instance, scaffold attachment factor B1 (SAFB1) is a large multi- functional protein involved in many cellular processes. Some of the major functions of SAFB1 are RNA processing and modulating cell growth (56). Experiments in mice showed that SAFB1 is essential for maintaining germinal tissue, suggesting an important role in reproduction (57). Succinate dehydrogenase B (SDHB) is upregulated in giant sea anemone tentacles indicating increased cellular respiration in these tissues. Transcriptional regulator ERG plays a major role in endothelial cell survival and interacts directly with YAP and the Hippo signaling pathway, influencing cell proliferation (58).

The giant sea anemones have an evolutionarily divergent expression pattern for genes involved in cell proliferation, metabolism regulation, and animal organ development. A shift in the expression of genes facilitating growth might be a specific evolutionary property of giant sea anemones, explaining how they can grow to their large sizes. Although our results suggest that the presence of anemonefish and *Symbiodiniaceae* affect gene expression evolution of certain groups of genes in giant sea anemone tentacles, we cannot conclusively determine how multi-organism symbiosis led to the evolution of specific gene expression patterns; viz our study cannot determine the mechanisms by which an ancestral gene expression pattern was co-opted to enable the multi-organism symbiosis. The divergent gene expression patterns in giant sea anemone tentacles could also be due to alternate developmental regimes. While our study cannot conclusively determine whether this lineage-specific gene expression regime is a direct consequence of the evolution of the multi-organism symbiosis, the adaptive benefits of the symbiosis in promoting the growth and survival of giant sea anemones (and, in turn, their evolution) are clear.

### The evolution of multispecies interaction promotes growth and survival

There is a strong link between the presence of anemonefish and nutrient utilization in sea anemone and their *Symbiodiniaceae*, suggesting that the symbiosis with anemonefish can have several adaptive advantages for both organisms (19, 24). The *Symbiodiniaceae* abundance in *Entacmaea quadricolor* that were reared with anemonefish was more than twice that in anemones without (36, 59). Nitrogen uptake by sea anemones with anemonefish was substantially lower than those without anemonefish, implying that *Symbiodiniaceae* in sea anemones hosting anemonefish were nitrogen sufficient (59). During periods of food scarcity, the sea anemone may not be able to meet the nitrogen demands of *Symbiodiniaceae*, thereby compromising their growth (60). Anemonefish provide an additional source of nitrogen that reduces the *Symbiodiniaceae’*s dependency on the host anemone feeding, thereby allowing even starved anemones to maintain high densities of *Symbiodiniaceae* (61).

In a long-term experiment, Holbrook and Schmitt observed that sea anemones with anemonefish grew more than three times as fast and were two-thirds bigger in size than anemones without anemonefish (62). The presence of anemonefish also enhanced reproductive activity in sea anemones, with hosting anemones asexually reproducing twice as fast as non-hosting anemones (62). Anemonefish can also enhance the growth of sea anemone and *Symbiodiniaceae* through increased oxygenation. Anemonefish exhibit several behaviors like rapid fin strokes and wiggling deeper into the anemone tentacles which improves water flow, thereby improving oxygen uptake by sea anemones, especially during periods of low photosynthetic activity at night (49).

These experimental results show that anemonefish are a vital nutrient source that can regulate metabolic processes in both *Symbiodiniaceae* and sea anemone; thus aligning with our observations of metabolic genes as being key targets in the evolution of symbiosis.

### Conclusion

We showed that in the tentacles of giant sea anemones, specific genes experienced divergent evolutionary shifts in expression. These expression shifts have clear benefits for the multispecies interaction between anemonefish and *Symbiodiniaceae* and could have evolved due to their symbiosis with anemonefish and *Symbiodiniaceae*. Using the insight gained from this study, we can propose a model by which the multi-organism symbiosis helps giant sea anemone attain their large sizes.

Juvenile sea anemones acquire *Symbiodiniaceae* from their environment. The nutrient exchange with *Symbiodiniaceae* promotes growth, allowing the anemone to reach sizes large enough to host anemonefish. Once anemonefish start occupying sea anemones, a tripartite relationship begins between anemonefish, sea anemone, and *Symbiodiniaceae*. The feeding and grooming behavior of anemonefish provide extra nutrients to sea anemone and *Symbiodiniaceae* increasing their metabolic output. Additionally, anemonefish protect their host anemones from predators like butterflyfish, ensuring that the giant sea anemones can grow to their large sizes.

Although the host anemones diversified before the origin of the symbiotic relationship with anemonefish, there is still a general effect of this relationship on tentacle gene expression in the host anemones. Furthermore, because of its early origin, the symbiosis with *Symbiodiniaceae* likely influenced gene expression in giant sea anemones longer than anemonefish. Disentangling the individual effects of anemonefish and *Symbiodiniaceae* on giant sea anemone gene expression will need targeted experiments. Using the functionally and evolutionarily important genes discovered in this study as candidates, future studies can help uncover specific mechanisms through which anemonefish and *Symbiodiniaceae* influenced the evolution of one of the most iconic symbioses in the animal kingdom.

## Materials and Methods

All code and supporting information can be found at https://github.com/agneeshbarua/Anemone_tentacle_gene_exp. Data generated in this study has been deposited in National Center for Biotechnology Information (NCBI) Sequence Read Archive (SRA) database under the BioProject PRJNA877849.

### Comparative transcriptomics

Expression data for giant sea anemone tentacles were collected from dissected tentacles of wild specimens found throughout southern Japan. Details about sampling locations, RNA extraction, and sequencing can be found in (22). All other data were obtained from the NCBI SRA database. Table S1 summarizes the species information and details of SRA id and bioproject.

SRA files were downloaded from NCBI using *prefetch* and *fasterq-dump* function of sratoolkit v2.10 (https://github.com/ncbi/sra-tools/wiki), followed by quality check using fastqc (https://www.bioinformatics.babraham.ac.uk/projects/fastqc/). All sequenced reads were processed using Trimmomatic v0.39 (66) to remove low-quality reads and Illumina adapters. Reads were assembled using Trinity, with redundant transcripts removed using a CD-HIT-EST cut-off of 95% (67). The initial assemblies contained transcripts from both cnidarians and *Symbiodiniaceae*. To remove these *Symbiodiniaceae* transcripts, we first calculated GC content for each transcript and separated the transcripts based on a GC content cut-off. Dinoflagellates and cnidarians have drastically different GC content (e.g. cnidarian ∼40% while *Symbiodiniaceae* <55%) (22); therefore, we used a GC content below 60% to filter out all cnidarian specific transcripts. Next, we created a BLAST dinoflagellate database with sequences from *Symbiodinium* (former clade A), *Breviolium* (former clade B) and *Cladocopium* (former clade C) and screened transcripts against this database using a BLASTN filter of 1e-20. The giant sea anemones had the highest number of hits to the dinoflagellate database, with hits primarily to *Cladocopium* (former clade C). This combined approach allowed us to separate the transcriptomes of cnidarians and any symbionts they might harbor.

After removing symbiont transcripts, corresponding proteins were predicted for each transcriptome using Transdecoder v5.5.0 (https://github.com/TransDecoder/TransDecoder). Only transcripts encoding an open reading frame of ≥100 were considered. Redundancy in sets of predicted proteins was further reduced by using a CD-HIT similarity cut-off of 95%. Lastly, since transcriptomes assembled by trinity represented all spice variants and thus multiple protein isoforms, we selected only the longest isoform as the representative isoform. The final list of predicted proteins matched previous estimates from other studies; for example, Putnam *et. al*. estimated *Nematostella vectensis* to contain ∼27,000 protein-coding genes, similar to our estimation of 26,698 protein-coding genes (68). The current gene set *Actinia tenebrosa* has ∼20,000 protein-coding genes, similar to our estimation of 21,197 (https://www.ncbi.nlm.nih.gov/data-hub/taxonomy/6105/). BUSCO v4.1.2 was used to check the completeness of the final using the final list of protein-coding genes (Table S2).

Quantification of libraries was done using kallisto v0.46.1 (69). Transcripts corresponding to the final list of protein-coding genes were used to create the kallisto index file. The final list of protein-coding genes were used as input for OrthoFinder v2.5.2 (OF) (70), with an MCL inflation parameter of 1.7 (higher stringency). OF assigned 87.2% of genes into 37,612 orthogroups, where 2496 orthogroups have sequences with all species present (Table S12). Using a custom R script we obtained a list of 1465 one-to-one orthologs expressed across all tissues in all taxa Table S13.

We used a new metric for quantifying RNA-seq reads called TPM10K (27). As the number of genes in a species varies, the transcript per million (TPM) values are not directly comparable (since the mean of libraries in TPM is 10^6^/*n*; where *n* = number of genes in reference). In a species with a higher degree of transcriptome completeness (i.e. higher number of genes *n*) all the genes would have a lower expression value. The TPM10K normalization accounts for this by multiplying the TPM value of a transcript by the number of genes in the reference assembly and dividing by an arbitrary number (in this case 104) to reduce the magnitude of expression value (27):

TPM10K*i* = TPM*i* x *n*/104 ; *i* = number of mapped reads to gene *i*.

Principal component analysis (PCA) was performed using the plotPCA function in *DESeq2*. Since we are comparing expression data from homologous tissues of multiple species from multiple studies, it is vital to remove their respective batch effects (71). To remove these batch effects and identify any existing patterns in expression between tissues and species, we used an empirical Bayes method implemented via the ComBat function in *sva* R package (72). This approach has been successfully implemented in previous studies (28, 73, 74).

### Analysis of phylogeny vs expression tree

Species phylogeny was constructed using 314 single-copy-orthologs estimated by OF. Multiple sequence alignment was done using MAFFT v7.3 (75). Phylogeny was constructed using IQTREE2 v2.1 (76) and ultrafast bootstrap was used to estimate node significance (77). Several amino acid models were tested using ModelFinder (within IQTREE), which determined JTT+F+R5 to be the best model based on Bayesian Information Criteria (BIC) (78). The tree was rooted at the Medusozoa and Anthozoa split using the reroot function in the R package *Phytools* (79).

Expression trees were constructed using mean tentacle gene expression from all samples species. The R package *ape* was used to construct a Neighbor-Joining expression tree based on 1-Spearman rank correlation distance. Branching patterns were assessed with bootstrap analysis using 1000 replicates.

To determine the relationship between expression distances and phylogenetic distances we first estimated pairwise distances between the pairs of tips from the phylogenetic tree and expression tree using the cophenetic.phylo function in the R package *ape* (80). Next, using these pairwise distance matrices, we performed a Mantel test using the R package *vegan* (81). The expression distance rates were calculated based on the slope of a linear regression between pairwise expression distances and phylogenetic distances.

### Co-expression and functional annotation

To get a sense of the biological underpinnings of the unique expression pattern of giant sea anemone tentacles, we first identified differentially expressed genes. Using the R package *edgeR*, we performed pairwise comparisons of *Heteractis, Stichodactyla, Macrodactyla*, and *Entacmaea* with *Actinia tenebrosa, Galaxea fascicularis, Nematostella vectensis*, and *Oulactis sp*. The comparisons were done one-to-one for each species of giant sea anemone and the rest, i.e. *Heteractis crispa* x *Actinia tenebrosa, Heteractis crispa* x *Galaxea fascicularis, Heteractis crispa* x *Nematostella vectensis, Heteractis crispa* x *Oulactis sp*, etc. Raw counts obtained from kallisto were converted to trimmed mean M values (TMM) for the DGE analysis. A complete list of differentially expressed genes can be found in Table S14. We next identified genes that were consistently upregulated and downregulated in all comparisons. Orthologs of all these genes in anemonefish hosting giant sea anemone were used as input to determine clusters of co-expressed genes using the R package *coseq* (82). *Coseq* performs clustering analysis of RNA-seq data using transformed profiles of raw counts. Based on the recommendation in the *coseq* vignette (https://www.bioconductor.org/packages/devel/bioc/vignettes/coseq/inst/doc/coseq.html), we implemented a Gaussian mixture model with arcsine transformation for its higher robustness. The clustering program was ran for a 100,000 iterations. *Coseq* includes genes into clusters using *Maximum a Posteriori* scores, a Bayesian optimized parameter search approach for estimating joint probability distributions.

Gene Ontology annotations were retrieved for each reference transcriptome using the PANNZER2 web server (83). We performed GO term enrichment using GOstas, GO.db, and GSEABase R packages (84–86). We modified the annotations to fit the format of the mentioned packages. A hypergeometric test with a false discovery rate (FDR) cut-off of 0.05 was used to consider a GO term as enriched.

### Phylogenetic ANOVA and Phylogenetic Multilevel Model

We carried out a phylogenetic analysis of variance (phy-ANOVA) using the Expression Variance and Evolution model developed by Rohlfs and Nielsen (35). The model describes phylogenetic expression level evolution between species and expression level variance within species. This approach can test for lineage-specific expression shifts in specific lineages (35). The test can be considered a phylogenetic analogy to tests for genetic drift via ratios of between and within species (population) variance. Expression data for orthologs in both up and down clusters and the species phylogeny were used to run the analysis.

The top ten orthologs that experienced the highest shifts in their expression throughout evolution (evolutionary divergent gene expression) were used for analysis using a phylogenetic multilevel model (PGMM). The PGMM was implemented using the *brms* R package (39). The rationale of the PGMM is similar to that of independent contrasts implemented through phylogenetic least squares, where the relationship between species is modelled to estimate the effect of evolutionary relationships on character evolution (87). In PGMM, the relationship between species is fit as a ‘group level’ effect (random effect in the terminology of mixed models). The relationship between species is fit using a phylogenetic covariance matrix (88). In our approach, we ran an intercept only model where the mean of the response variable (gene expression of ten genes) is conditioned across group level effects of phylogeny, presence of anemonefish, and presence of *Symbiodiniaceae*. In other words, we estimate whether differences in gene expression between species are influenced by the presence of anemonefish and *Symbiodiniaceae* while accounting for the evolutionary relationships of the different cnidarian species.

We used a pseudoalignment approach to determine the *Symbiodiniaceae* content in our species. Using transcriptomes of C1 clade such as *Symbiodinium aenigmaticum, Symbiodinium minutum, Symbiodinium pseudominutum, Symbiodinium psygomophilum*, and clade C1 associated *Symbiodinium* from *Siderastrea siderea*, (see Table S15 for links to files) we build a composite *Symbiodiniaceae* transcriptome that we used an input to build a kallisto index. We then used this index to quantify reads matching to *Symbiodiniaceae* in the SRA fastq files (not the *de novo* assembled transcriptomes). We then estimated the total number of reads pseudoaligned to the composite *Symbiodiniaceae* transcriptome (Fig S7). In our modelling approach, we used the quantification of *Symbiodiniaceae* composition to help us classify cnidarian species as having or not having *Symbiodiniaceae*.

We used the *get_prior* function in the *brms* package to get standard priors and fit the model as a ‘gaussian’ distribution of the response variable. The model was run for 5000 iterations with an initial burin of 3000. The effective sample size from the posterior was high and convergence of the MCMC was judged using the *Rhat* value, where an *Rhat* value considerable greater than 1 (i.e., > 1.1) indicated chains have not converged. The *Rhat* values of our estimates ranged from 1.0 to 1.01, indicating convergence of chains.

The top 10 genes were annotated using a combination using of blastp and Uniprot. We used the one-to-one orthologs of the top 10 genes in *Nematostella vectensis* as query sequences for blastp and selected maximum 5 sequences that had an e-value lower than 1e-10. Using the blastp output, we obtained annotations from UniProt. Only ‘reviewed’ annotations were kept. The final annotations are available in Table S11.

## Acknowledgments

The authors would like to thank the OIST Sequencing Center (SQC) for their support during sequencing of the samples.

## Notes

### Competing Interest Statement

The authors have declared no competing interest.

