## Supplementary_figures for "The genetic basis for adaptation in giant sea anemones to their symbiosis with anemonefish and *Symbiodiniaceae*"

Agneesh Barua

August 2022

#### Figure S1: Spearmans correlations for all species and all tissues

The gene expression of jellyfish tentacles was highly distinct, with very low similarities between jellyfish and non-jellyfish samples

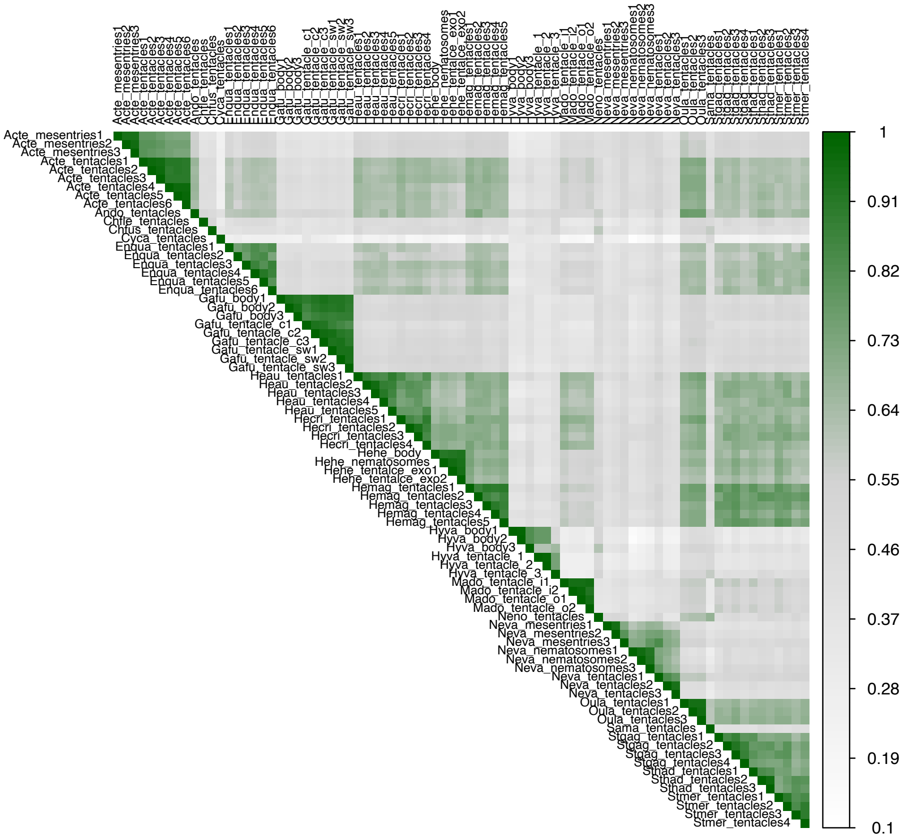

**Figure S2: Spearmans correlations for non-actinaria and Heteractis-Stichodactyla**

The following figure contrasts the difference in expression between jellyfish and giant sea anemones *Heteractis* and *Stichodactyla*

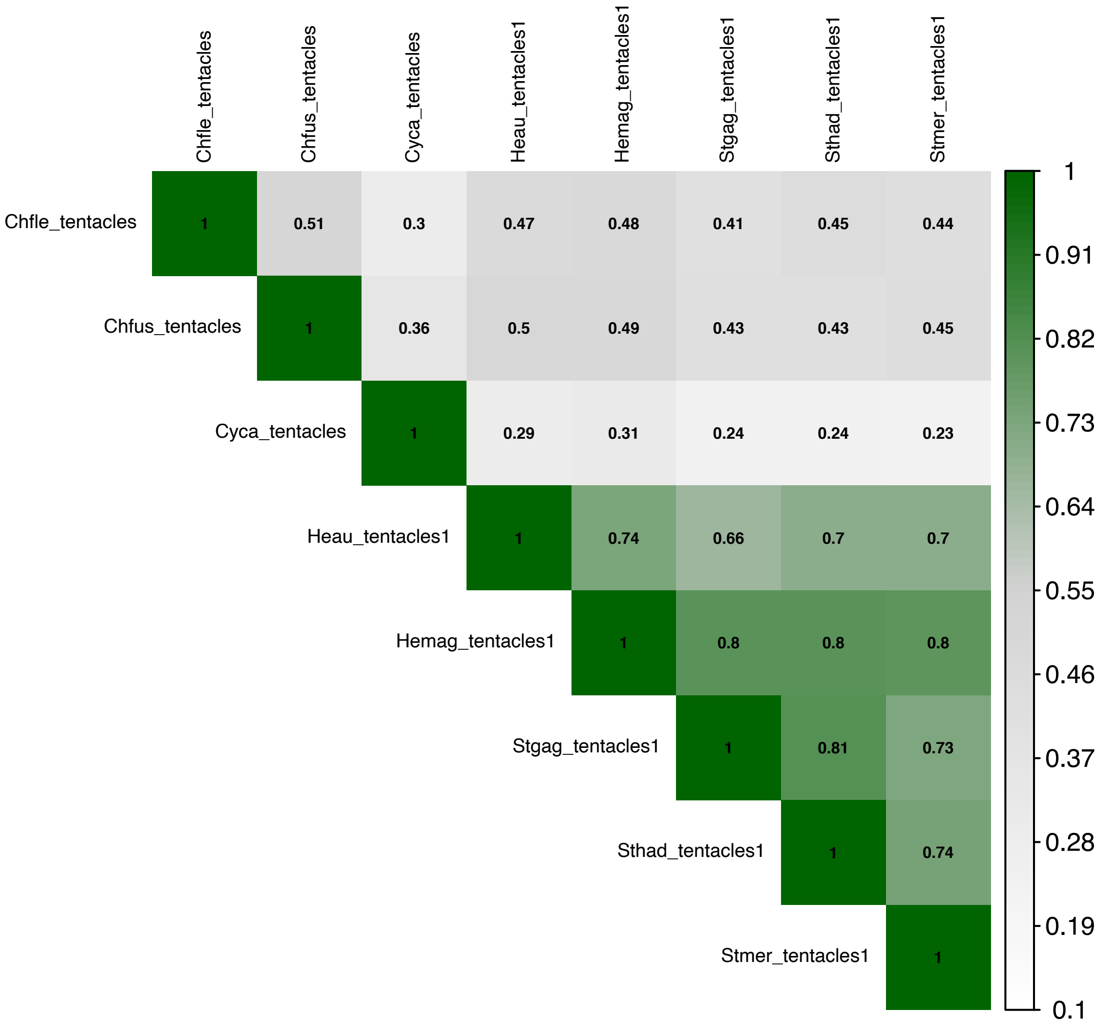

### Figure S3: Spearmans correlations for all actinaria tentacles

Gene expression within actinaria was overall more similar than between actinaria and non-actinaria

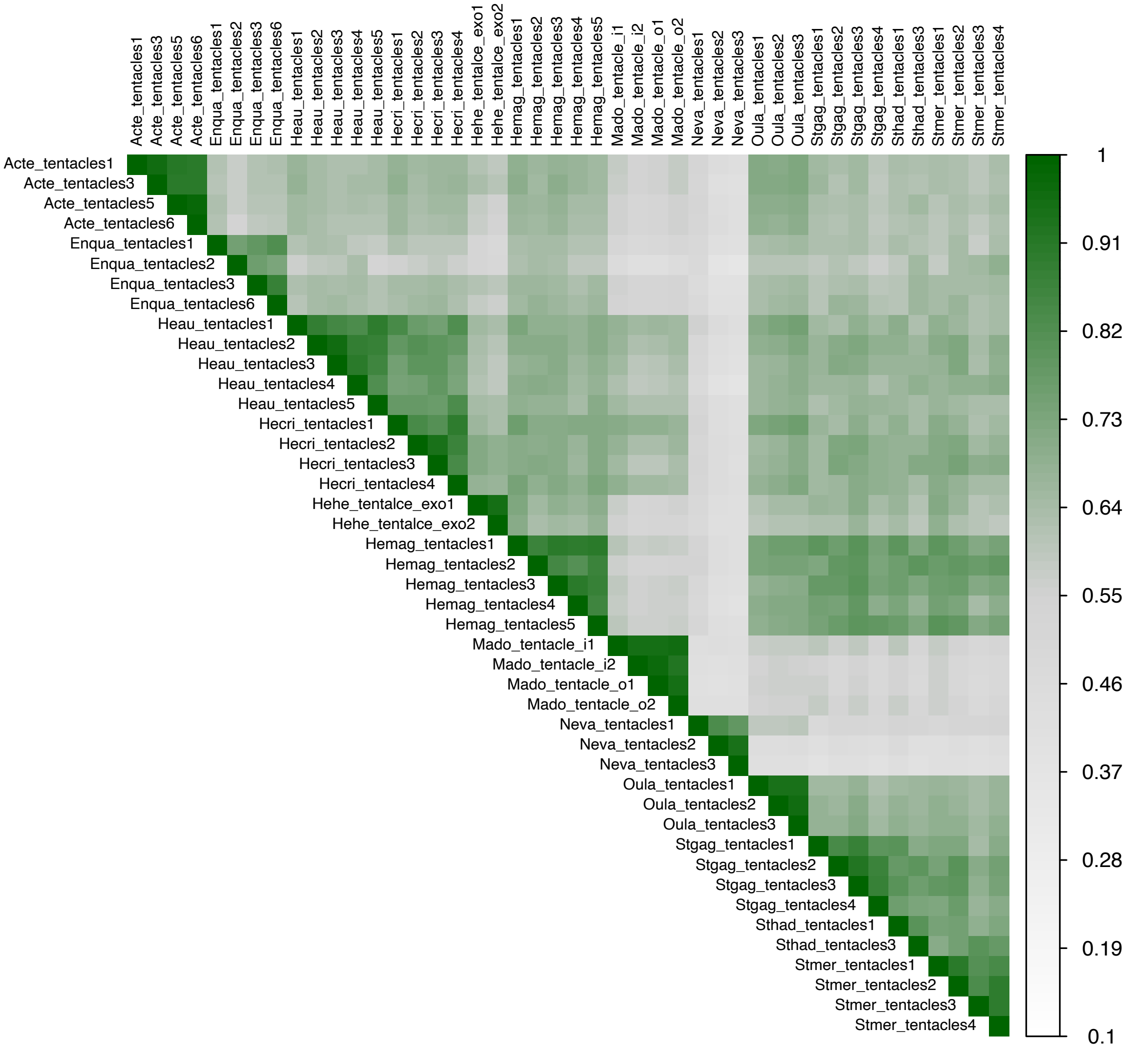

### Figure S4: PCA for all cnidaria samples

The PCA with all cnidarian samples is included in the supplementary information and showed the same overall trends as ones with only anthozoans.
The first two components partition the data based on tissues.

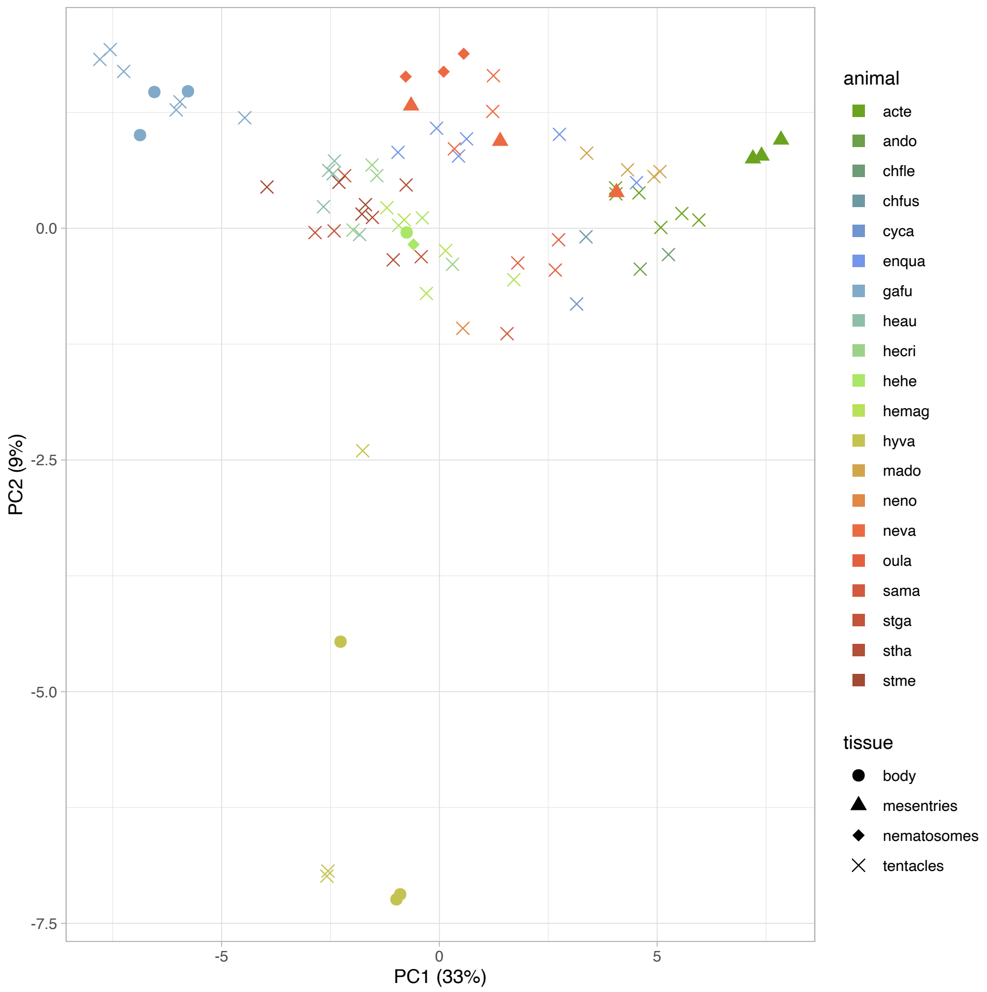

**Figure S5: Spearmans correlations for random set of 47 genes**

We estimated correlations using a random assortment of 47 genes to check if this was the case. A random set of 47 genes showed less noisy correlation patterns (Fig S5), suggesting that the 47 genes in cluster 1 are highly variable due to biological factors and not sampling bias.

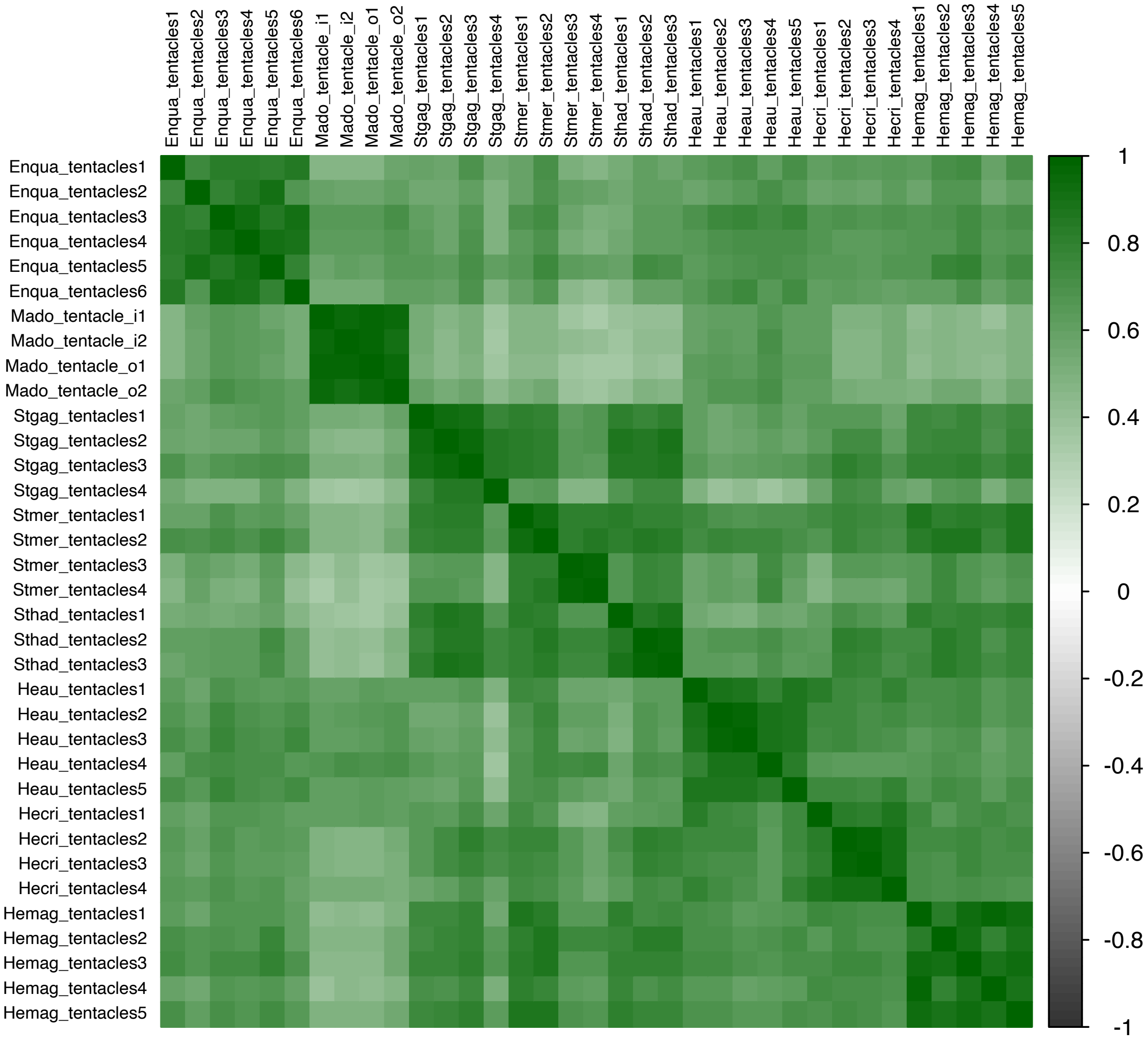

### Figure S6: Coseq analysis for downregulated orthologs

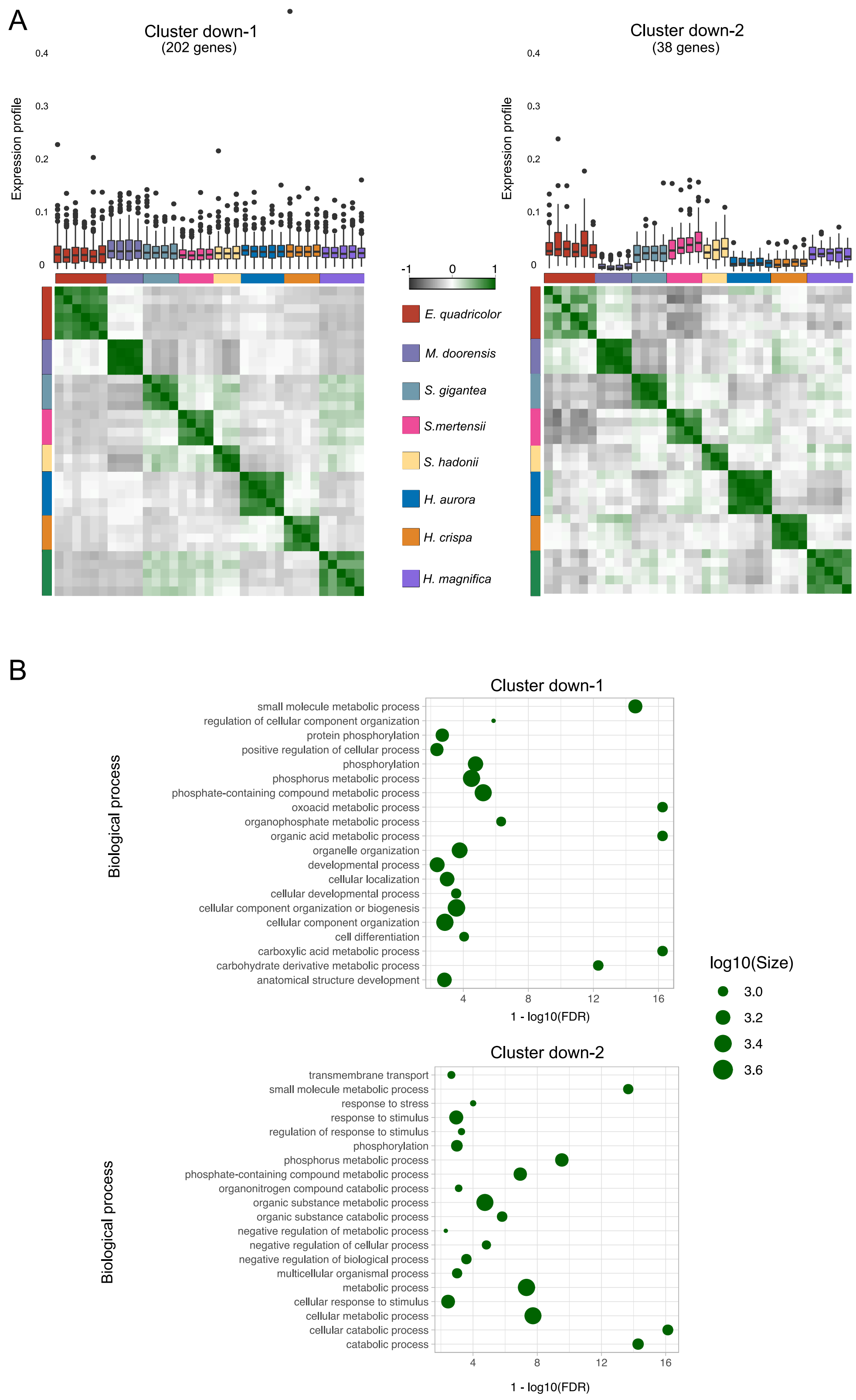

### Figure S7: Symbiodiniaceae abundance

We used a pseudoalignment approach to determine the Symbiodiniaceae content in our species. The following figure shows the total number of reads pseudoaligned to the composite Symbiodiniaceae transcriptome.

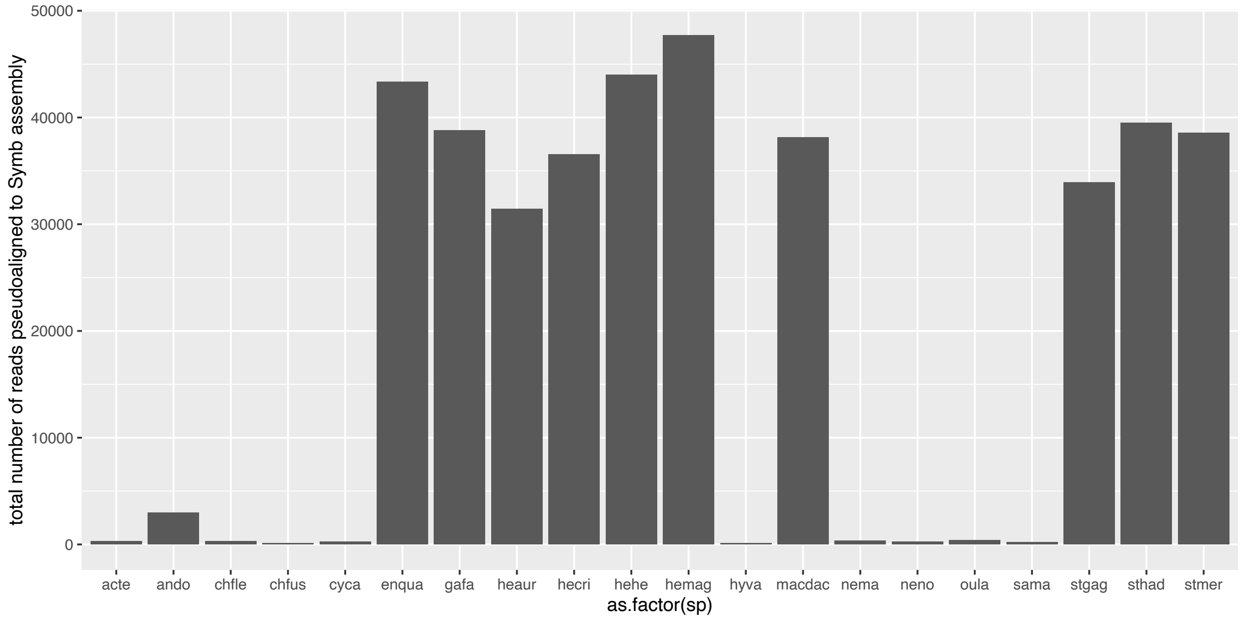
